# Mck1 defines a key S-phase checkpoint effector in response to various degrees of replication threats

**DOI:** 10.1101/610105

**Authors:** Xiaoli Li, Xuejiao Jin, Sushma Sharma, Xiaojing Liu, Jiaxin Zhang, Yanling Niu, Jiani Li, Zhen Li, Jingjing Zhang, Qinhong Cao, Beidong Liu, Huiqiang Lou

**Author notes:** Department of Pathology, UT Southwestern Medical Center, Dallas, TX 75390-9072, USA. Department of Molecular and Human Genetics, Baylor College of Medicine, Houston, TX 77030, USA.

## Abstract

The S-phase checkpoint plays an essential role in regulation of the ribonucleotide reductase (RNR) activity to maintain the dNTP pools. How eukaryotic cells respond appropriately to different levels of replication threats remains elusive. Here, we have identified that a conserved GSK-3 kinase Mck1 cooperates with Dun1 in regulating this process. Deleting *MCK1* sensitizes *dun1*Δ to hydroxyurea (HU) reminiscent of *mec1*Δ or *rad53*Δ. As a kinase at the downstream of Rad53, Mck1 does not participate in the post-translational regulation of RNR as Dun1 does, but Mck1 can release the Crt1 repressor from the promoters of *RNR2/3/4* by phosphorylation. Meanwhile, Hug1, an Rnr2 inhibitor, is induced to fine-tune the dNTP levels. When cells suffer a more severe threat, Mck1 can inhibit the transcription of *HUG1*. Importantly, only a combined deletion of *HUG1* and *CRT1*, can confer a dramatic boost of dNTP levels and the survival of *mck1*Δ*dun1*Δ or *mec1*Δ cells assaulted by a lethal dose of HU. These findings reveal the division-of-labor between Mck1 and Dun1 at the S-phase checkpoint pathway to fine-tune dNTP homeostasis.

**Author Summary:** The appropriate amount and balance of four dNTPs are crucial for all cells correctly copying and passing on their genetic material generation by generation. Eukaryotes have developed an alert and response system to deal with the disturbance. Here, we uncovered a second-level effector branch. It is activated by the upstream surveillance kinase cascade, which can induce the expression of dNTP-producing enzymes. It can also reduce the inhibitor of these enzymes to further boost their activity according to the degrees of threats. These findings suggest a multi-level response system to guarantee dNTP supply, which is essential to maintain genetic stability under various environmental challenges.

## Introduction

To ensure the genome stability, the DNA replication process is under strict surveillance by the S-phase checkpoint (also known as the intra-S or replication checkpoint) in all eukaryotes [1–5]. The main kinases of the cascade, Mec1^ATR^ and Rad53^CHK2^, are activated in response to aberrations in DNA replication [6–10]. Among all the various downstream effects, the essential role of Mec1-Rad53 has been demonstrated to be the regulation of the activity of RNR in *Saccharomyces cerevisiae* [3, 11, 12].

RNR catalyzes the reduction of ribonucleoside diphosphates to their deoxy forms, which is the rate-limiting step in the *de novo* synthesis of deoxyribonucleoside triphosphates (dNTPs), the building blocks of DNA [13]. RNR is normally composed of two large subunits R1 (Rnr1 homodimer) and two small subunits R2 (Rnr2 and Rnr4 heterodimer) in budding yeast. Proper and balanced cellular dNTP pools are essential for genome integrity [14, 15]. Therefore, several RNR inhibitors such as hydroxyurea (HU), clofarabine and gemcitabine have been exploited for the chemotherapy of several types of cancers [16]. The RNR activity is strictly controlled by multi-layer mechanisms in cells [15, 17]. First, RNR is allosterically regulated through the binding of different forms of effector nucleotides, for example, ATP or dATP. Second, the expression of *RNR1-4* genes is controlled at both transcriptional and post-transcriptional levels. For instance, *RNR1* gene is activated during G_1_/S transition by the MBF transcription factor, whilst the excessive expression of *RNR2-4* is repressed by Crt1 (<u>c</u>onstitutive <u>R</u>NR transcription 1) through recruiting the Ssn6-Tup1 co-repressor complex to the promoter. Furtheremore, *RNR3*, as a *RNR1* paralog, is generally silenced till the release of Crt1 under stressed condition [18]. Third, the RNR enzyme activity is post-translationally inhibited by several small intrinsically disordered proteins such as Sml1, Dif1 and Hug1 in *S. cerevisiae* and Spd1 in *Schizosaccharomyces pombe* [15]. Sml1 binds to cytosolic Rnr1 and disrupts the regeneration of the Rnr1 catalytic site [19, 20]. Dif1 promotes the nuclear import of the Rnr2/Rnr4 heterodimer, which is anchored by Wtm1 in the nucleus [21–23], precluding Rnr2/Rnr4 from associating with Rnr1 or Rnr3 to form the RNR holo-enzyme in the cytoplasm. Hug1, like Dif1, also contains a HUG domain, which can inhibit RNR through binding Rnr2 [24, 25].

When cells encounter genotoxic agents, RNR is stimulated by the Mec1-Rad53-Dun1 kinase cascade at both transcriptional and post-translational levels to provide adequate dNTPs for DNA replication/repair [13, 15, 26–29]. One of the key Mec1-Rad53-Dun1 targets is Crt1, which becomes hyperphosphorylated and therefore leaves the promoter of damage-inducible genes such as *RNR2-4*, *HUG1* and *CRT1* itself [18]. Apart from the Crt1 repressor, Dun1 also targets RNRs’ protein inhibitors including Sml1 and Dif1 [21–23, 30, 31], both of which are hyperphosphorylated and degraded [22, 32]. Spd1 is degraded in S phase and after DNA damage via the ubiquitin-proteasome pathway as well [33]. Unlike Sml1 and Dif1, Hug1 is induced together with Rnr2-4 due to the removal of the Crt1 repression from its promoter in the presence of genotoxic agents. As a result, Hug1 acts in a distinct but undefined manner compared with its paralogs Sml1 and Dif1 [25]. Intriguingly, the lethality of *mec1*Δ or *rad53*Δ can be suppressed by deleting any of the negative regulators of RNR mentioned above (*CRT1*, *SML1*, *DIF1* or *HUG1*) [9, 12, 34, 35]. All these findings highlight the importance of RNR regulation by Mec1-Rad53-Dun1.

In this study, we identify that a combinational deletion of *MCK1* and *DUN1* displays a synergistic effect, reminiscent of the extreme sensitivity of *mec1*Δ or *rad53*Δ to HU. No effects are observed when we delete other glycogen synthase kinase-3 (GSK-3) homologs such as *YGK3*, *MRK1* and *RIM11* in *S. cerevisiae*. Also, Rad53 kinase is able to phosphorylate Mck1 *in vitro*. Moreover, deletion of *CRT1* suppresses the HU sensitivity of *dun1*Δ*mck1*Δ. Crt1 phosphorylation is significantly compromised in *mck1*Δ accompanied with dissociation of Crt1 from the *RNR* promoters and reduction of inducible *RNR3* expression. Apart from Crt1, Mck1 also negatively regulates the *HUG1* transcription. Taken together with previous findings, these data suggest that Mck1 and Dun1 define two non-redundant and cooperative branches of the Mec1-Rad53 kinase cascade in fine-tuning RNR activity when cells encounter replication stress.

## Results

### Mck1 plays a vital role in coping with replication stress in the absence of Dun1

Deletion of *SML1*, encoding an Rnr1 inhibitor, is known to suppress the lethality of *mec1*Δ or *rad53*Δ cells [34] (Fig. 1A). Nevertheless, *mec1*Δ*sml1*Δ or *rad53*Δ*sml1*Δ are extremely sensitive to HU (Fig. 1B). On the other hand, deletion of the only known downstream kinase of Mec1-Rad53 [27], *DUN1*, resulted in a much lower HU sensitivity than that of *mec1*Δ*sml1*Δ or *rad53*Δ*sml1*Δ. These data raise the possibility that there might be Dun1-independent players downstream the Mec1-Rad53 pathway working in parallel with Dun1 in response to the RNR inhibitor [18, 27] (Fig. 1A).

**Figure 1.**
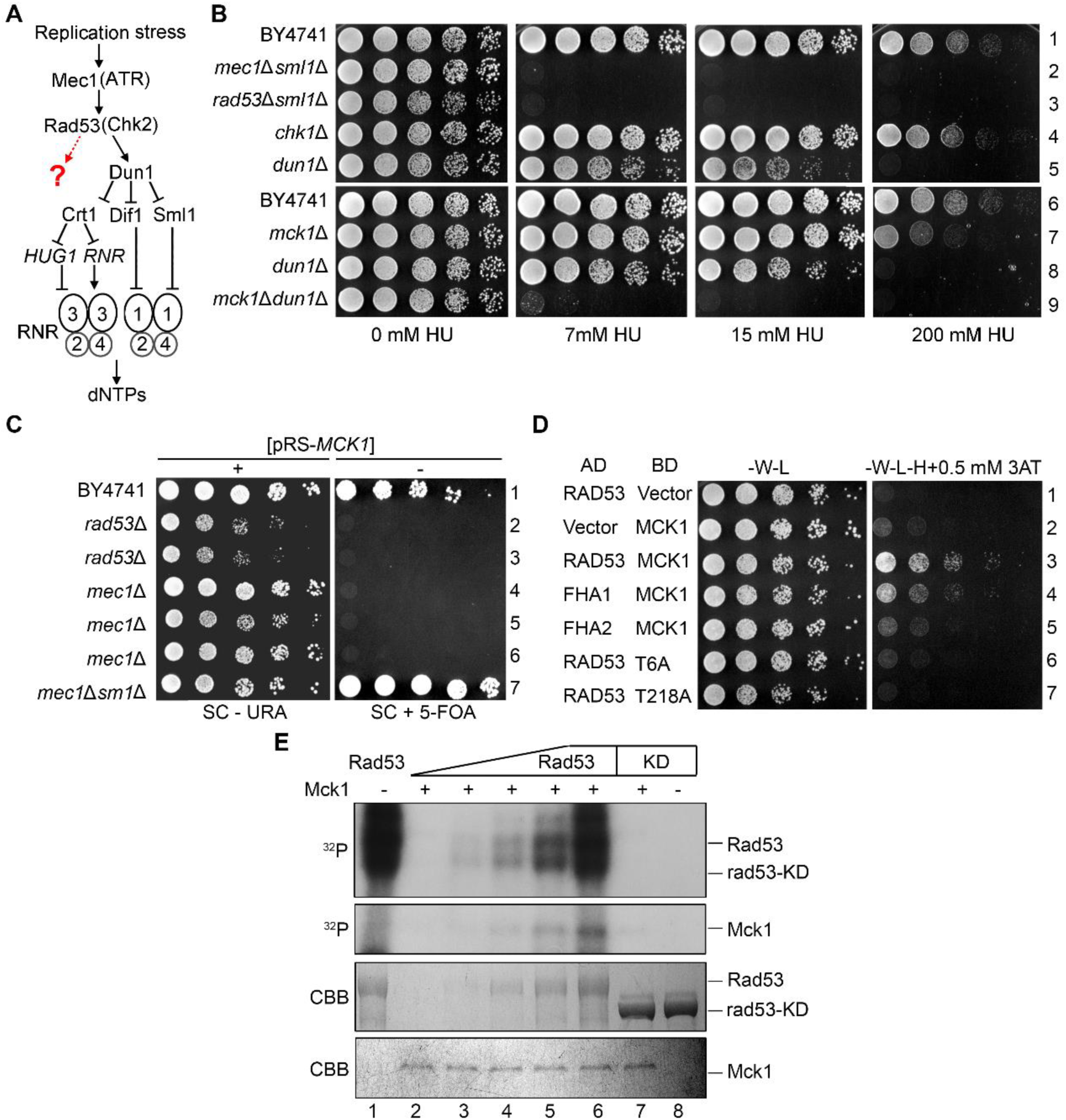
*MCK1* is indispensable for *dun1*Δ cells dealing with replication stress. (A) Regulation of celluar dNTP levels by the S phase checkpoint in *S. cerevisiae*. (B) 5-fold serial dilutions of WT*, rad53*Δ*sml1*Δ*, mec1*Δ*sml1*Δ*, chk1*Δ*, dun1*Δ, *mck1*Δ, *dun1*Δ*mck1*Δ (Table S1) cells were spotted onto a YPD plate supplemented with the indicated concentrations of HU. Plates were incubated at 30°C for 48h before photograph. (C) Overexpression of *MCK1* rescues the lethality of *mec1*Δ *or rad53*Δ. 5-fold serial dilutions of the indicated strains were spotted onto SC-URA or SC+5FOA (5-fluoroorotic acid) plates and grown at 30°C for 48h. The plasmid expressing WT *MCK1*/*URA3* was lost in the presence of 5-FOA. (D) Physical interaction between Mck1 and Rad53. *MCK1* and *RAD53* were cloned into pGADT7 and pGBKT7 vectors, respectively. AH109 strains transformed with the indicated plasmids were grown on the SC-Trp-Leu or SC-Trp-Leu-His plates. 0.5 mM 3-amino-triazole (3AT) was added to inhibit the leaky *HIS3* expression. (E) Rad53 phosphorylates Mck1 *in vitro*. Mck1-5FLAG was precipitated from yeast cells and incubated with purified recombinant His6-RAD53 and His6-rad53-KD (K227A) in the presence of ɣ-^32^P-ATP as detailed in Methods. After resolved in an 8% polyacrylamide gel with SDS, the samples were subjected to autoradiography. Then, the gel was stained by Coomassie Brilliant Blue (CBB) to show the amount of the loaded protein in each reaction.

We identified candidates of the novel Mec1-Rad53 downstream players using the synthetic genetic array approach [36, 37]. A *DUN1* null mutant was crossed with the single deletion library. After acquiring the final double mutants through pinning on a series of selective media, the growth of the double mutants was analyzed in the presence or absence of HU. Gene mutants, when in combination with the *DUN1* deletion, that showed a synthetically sick or lethal phenotype under the HU stress condition were selected as the potential candidates of the Mec1-Rad53 downstream players. Among them, the *mck1*Δ*dun1*Δ double mutant grew normally in the absence of HU (Fig. 1B). However, it showed a remarkable HU sensitivity similar to that of *rad53*Δ*sml1*Δ or *mec1*Δ*sml1*Δ. The *mck1*Δ single mutant alone displayed mild sensitivity to 200 mM HU. These results indicate that Mck1 might work in parallel with Dun1 and has a Dun1-independent function in the cell’s survival under the HU replication stress.

Given that *MCK1* encodes one of the GSK-3 family serine/threonine kinases in budding yeast, we deleted its paralog *YGK3* and orthologs *MRK1* and *RIM11*. None of them exhibited HU sensitivity (Fig. S1A) or synthetic interaction with *DUN1* (Fig. S1B), indicating that GSK-3 kinase Mck1 might have a specific role in replication checkpoint, and this function is not shared by its paralogs. Meanwhile, GSK-3 kinases regulate transcription of the general stress responsive genes (e.g., glucose starvation, oxidative, heat shock and low pH) through two partially redundant transcription activators Msn2 and Msn4 [38]. However, deletion of both *MSN2* and *MSN4* showed no additive effect with *dun1*Δ (Fig. S1B), implying that Msn2/Msn4 is unlikely the major effectors of Mck1 in response to HU.

In addition to *DUN1*, *MCK1* showed genetic interactions with checkpoint activators and mediators as well. Deletion of *MCK1* markedly exacerbated the HU sensitivity of *mre11*Δ, *ddc1*Δ, *mrc1*Δ or *rad9*Δ (Fig. S2A and S2B), further arguing for a critical role of Mck1 in the S-phase checkpoint pathway.

### Mck1 is a downstream target of Rad53

The synergistic HU sensitivity caused by the combined deletion of *MCK1* and *DUN1* raises a possibility that they may function cooperatively in the Mec1-Rad53 pathway. We first tested whether *MCK1* is a dosage suppressor of the *mec1*Δ or *rad53*Δ lethality. We constructed a high-copy number plasmid with a *URA3* marker (pRS426) expressing *MCK1* and introduced it into diploid strains wherein one copy of *MEC1*, *RAD53* and *SML1* was deleted. After sporulation, the tetrads were analyzed by microscopic dissection. *mec1*Δ and *rad53*Δ spores could hardly grow unless carrying the pRS426-*MCK1* plasmid or in the absence of *SML1* (Fig. S2C). To verify it, we induced loss of this plasmid on a plate containing 5-floroorotic acid (5-FOA). Without the *MCK1* overexpression plasmids, neither *mec1*Δ nor *rad53*Δ was able to survive (Fig. 1C). These results indicate that *MCK1* overexpression is able to bypass the essential function of *MEC1* and *RAD53*, validating the results of previous large scale screen [39].

We then examined whether there is physical interaction between Mck1 and Mec1 or Rad53 using yeast two-hybrid assay and found that Mck1 shows positive interaction with Rad53 (Fig. 1D). To determine which part of Rad53 is required for the interaction with Mck1, we expressed the forkhead homology-associated domains (FHA1 and FHA2) of Rad53 protein, which are known to mediate interaction with other proteins. We found that FHA1 domain is sufficient to interact with Mck1. FHA1 preferentially binds the phosphothreonine (pThr) peptides bearing a pThr-x-x-D/E/I/L motif (x stands for any amino acid) [29]. Therefore, we mutated all six threonine residues to alanines in Mck1 (mck1-T6A), resulting in abolished interaction with Rad53 (Fig. 1D). Among these six threonine residues, the T218A mutation dramatically reduced the Mck1-Rad53 interaction. These results suggest that Mck1 interacts with the FHA1 domain of Rad53 through a canonical phosphorylation-mediated mechanism.

Given the physical association of Mck1 and Rad53, we tested whether Mck1 is a substrate of Rad53. We expressed and purified Rad53 or rad53-KD (a kinase-dead mutant, rad53-K227A) for *in vitro* kinase assays. Rad53 wild-type (WT), but not the KD mutant, showed robust auto-phosphorylation as indicated by the incorporation of ^32^P and by the electrophoretic shift (Fig. 1E, upper panel, compare lane 1 with 8), indicating that the robust kinase activity is Rad53-specific. Next, we isolated the endogenous FLAG-tagged Mck1 as a substrate through immunoprecipitation via anti-FLAG beads followed by FLAG peptide elution. With the increasing amounts of Rad53 kinase added in the reactions, more ^32^P was transferred to Mck1 (Fig. 1E, middle panel, lanes 2-6). On the other hand, rad53-KD completely failed to phosphorylate the Mck1 substrate (lane 7). A Coomassie brilliant blue (CBB) stained gel revealed nearly equal loading of Mck1 substrates in each reaction (Fig. 1F, lower panel). These results suggest that Rad53 kinase is able to phosphorylate Mck1 *in vitro*. Taken together with the synthetic genetic interaction between *MCK1* and *DUN1*, these data also suggest that Mck1 defines a new downstream branch of Rad53 in parallel with Dun1.

### Mck1 does not function through RNR sequesters including Sml1, Dif1 and Wtm1

To investigate the exact role of Mck1 in the Mec1-Rad53 pathway, we first asked whether *SML1* is a potent suppressor of *mck1*Δ as well. Surprisingly, deletion of *SML1* was not able to suppress the checkpoint defect in *mck1*Δ, in stark contrast to its capability to bypass the essentiality of *MEC1* or *RAD53* (Fig. 2A). Consistently, in the absence of Mck1, Sml1 was not affected at either mRNA or protein level upon HU treatment compared to WT (Fig. 2B and 2C). Interestingly, other known effectors of Dun1, e.g., Dif1 and Wtm1, were not the suppressors of *mck1*Δ as well (Fig. 2A). These results suggest that Sml1/Dif1/Wtm1 are not the downstream effector of Mck1, which are mainly targeted by the Dun1 branch.

**Figure 2.**
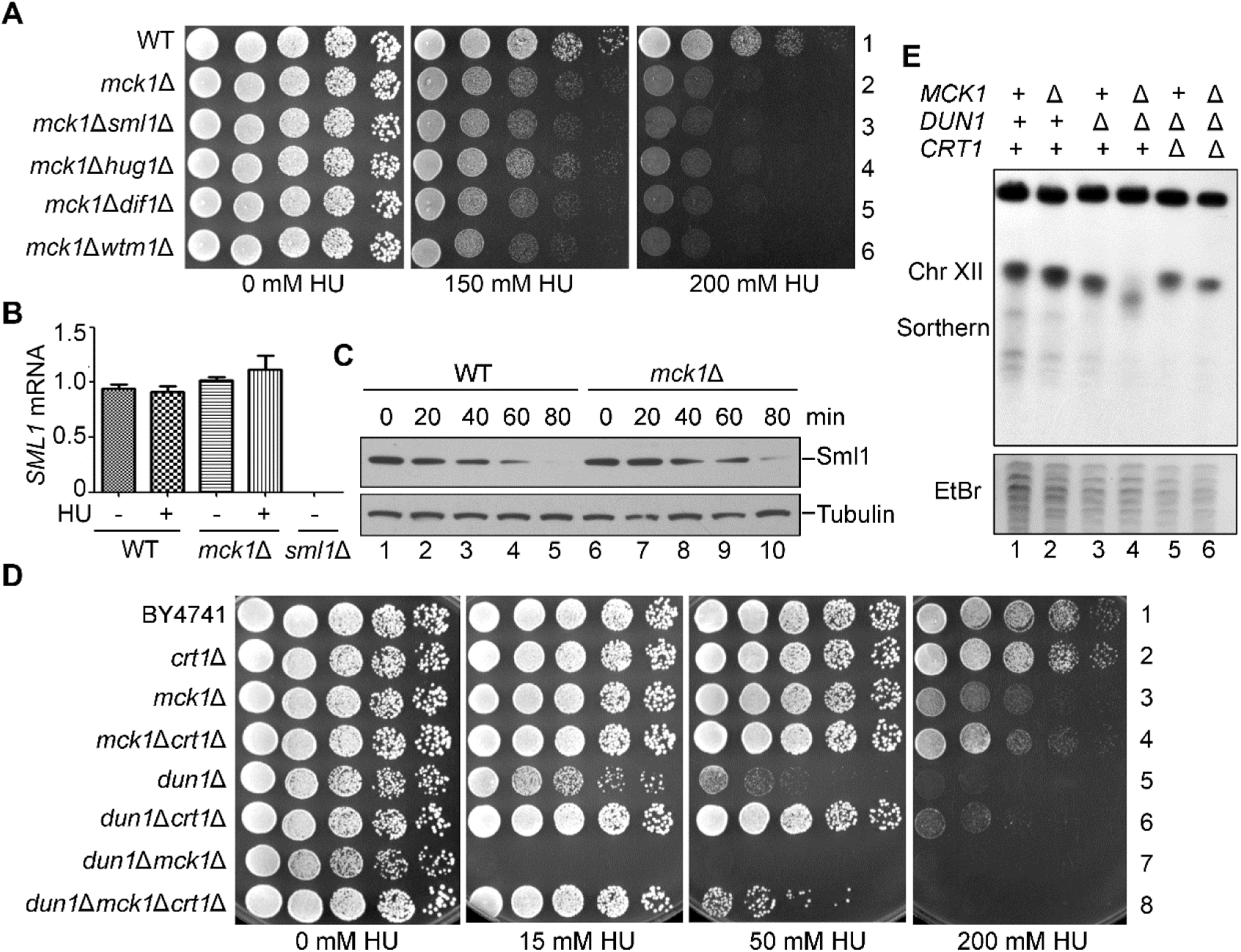
*crt1*Δ, but not *sml1*Δ, suppresses the checkpoint deficiency of *mck1*Δ. (A) Deletion of *SML1*, *DIF1* or *WTM1* has no apparent suppression of the HU sensitivity of *mck1*Δ. WT, *mck1*Δ, *mck1*Δ*sml1*Δ, *mck1*Δ*dif1*Δ, *mck1*Δ*wtm1*Δ (Table S1) were tested for the growth in the presence of the indicated concentrations of HU by serial dilution analysis as described in Fig.1A. (B) Measurement of the *SML1* mRNA levels in *mck1*Δ by qPCR. Cells were grown in rich media with or without 200 mM HU for 3h. The total mRNA was prepared as described in Methods and Materials. The relative levels of *SML1* to *ACT1* mRNAs were determined by qPCR analyses. Error bars represent standard deviations from at least three biological repeats. No significance was found by student *t*—test. (C) Mck1 is not involved in Sml1 degradation. The Sml1 protein levels in *mck1*Δ were examined after 200 mM HU treatment for the indicated times. YFP-Sml1 and Tubulin were detected by immunoblots. (D) Deletion of *CRT1* partially suppresses the HU sensitivity of *mck1*Δ *and mck1*Δ*dun1*Δ. WT, *crt1*Δ*, mck1*Δ*, mck1*Δ*crt1*Δ*, dun1*Δ*, dun1*Δ*crt1*Δ*, dun1*Δ*mck1*Δ*, dun1*Δ*mck1*Δ*crt1*Δ (Table S1) were tested for the HU sensitivity by serial dilution analysis as described in (A). (E) Mck1 and Dun1 play non-redundant roles in the rDNA copy number maintenance. Genomic DNA was prepared in an agar plug and separated on a 1% pulsed field gel. Ethidium bromide staining of yeast chromosomes is shown in the lower panel. Upper panel shows chromosome XII by hybridization with a probe against 18S rDNA.

### Mck1 regulates genome stability in a Dun1-independent manner

Besides these RNR sequesters, RNR is also controlled at the transcription level mainly through the repressor Crt1 (also called Rfx1) [18]. Indeed, deletion of *CRT1* significantly suppressed the sensitivity of *mck1*Δ and *dun1*Δ mutants to 200 mM and 50 mM HU, respectively (Fig. 2D). Intriguingly, deletion of *CRT1* conferred a better growth of *mck1*Δ*dun1*Δ double mutant than that of *dun1*Δ in the presence of up to 50 mM HU (compare lines 5 and 8). These results suggest that Crt1 is controlled by Dun1 and Mck1 in a non-redundant manner. Mck1 is required for efficient *RNR* induction through antagonizing Crt1, particularly in the presence of high concentration of HU (e.g., 200 mM).

The low RNR level causes insufficient dNTP supply, which leads to genome instability such as the copy number change of ribosomal DNA (rDNA) located at chromosome XII in *S. cerevisiae* [40]. Therefore, we examined the rDNA copy number by pulsed-field gel electrophoresis (PFGE) followed by Southern blotting. Without HU treatment, *mck1*Δ exhibited an rDNA copy number loss phenotype less than *dun1*Δ (Fig. 2E, compare lanes 2 and 3). However, the *mck1*Δ*dun1*Δ double mutant showed a more severe rDNA repeat loss than each single mutant (lane 4). Moreover, deletion of *CRT1* prominently ameliorated the rDNA instability phenotype of *dun1*Δ*mck1*Δ (lane 6). The degrees of rDNA copy number loss in these mutants correlated well with their growth defects in the presence of HU (compare Fig. 2D and 2E). These data suggest that Mck1 and Dun1 contribute independently to rDNA/genome stability through regulating Crt1 and thus *RNR* expression.

### Mck1 is involved in Crt1 phosphorylation

To address whether the kinase activity of Mck1 is required for *RNR* regulation, we mutated two conserved residues within the catalytic core (D164) and activation-loop (Y199) of Mck1 which have been demonstrated to be indispensable for its kinase activity [41]. Both *mck1-D164A* and *mck1-Y199F* showed synthetic lethality with *dun1*Δ in the presence of 50 mM HU (Fig. S3A), demonstrating that Mck1’s kinase activity is indispensable in response to replication stress.

It is known that the repression function of Crt1 is relieved through phosphorylation by Dun1 [18]. Since *CRT1* is a common suppressor for both *mck1*Δ and *dun1*Δ as mentioned above, we then hypothesized that Mck1 kinase may function through Crt1 phosphorylation as Dun1. To test this, we assessed the Crt1 phosphorylation levels through western blotting. As reported previously [18], Crt1 displayed a slower mobility shift (Crt1-P) after separation in a high resolution polyacrylamide gel (Fig. S3B). There was a basal level of Crt1 phosphorylation, which was largely dependent on Mck1 (Compare lanes 2-4). The Crt1 phosphorylation level increased significantly following 200 mM HU treatment for 3 h in WT. Deletion of *RAD53* or *MCK1* caused a relatively lower level of Crt1 phosphorylation than *DUN1* deletion, indicating the contribution of the Rad53-Mck1 branch in targeting Crt1.

Because Crt1 phosphorylation is cell-cycle-regulated, we next examined its level in the synchronized cell samples. Cells were synchronized by α-factor in G_1_ and released into the fresh media for the indicated time. Cell cycle progression was monitored by fluorescence-activated cell sorting (FACS). Under normal condition, Crt1 occurred at the beginning of S phase and reached a peak at the end of S phase (60 min) in WT (Fig. 3A, 3B and S3C). *MCK1* deletion caused a decrease in Crt1 phosphorylation, whereas combined deletion of *MCK1* and *DUN1* nearly abolished Crt1 phosphorylation. These results allow us to conclude that Mck1 and Dun1 function non-redundantly in Crt1 phosphorylation during normal S phase progression.

**Figure 3.**
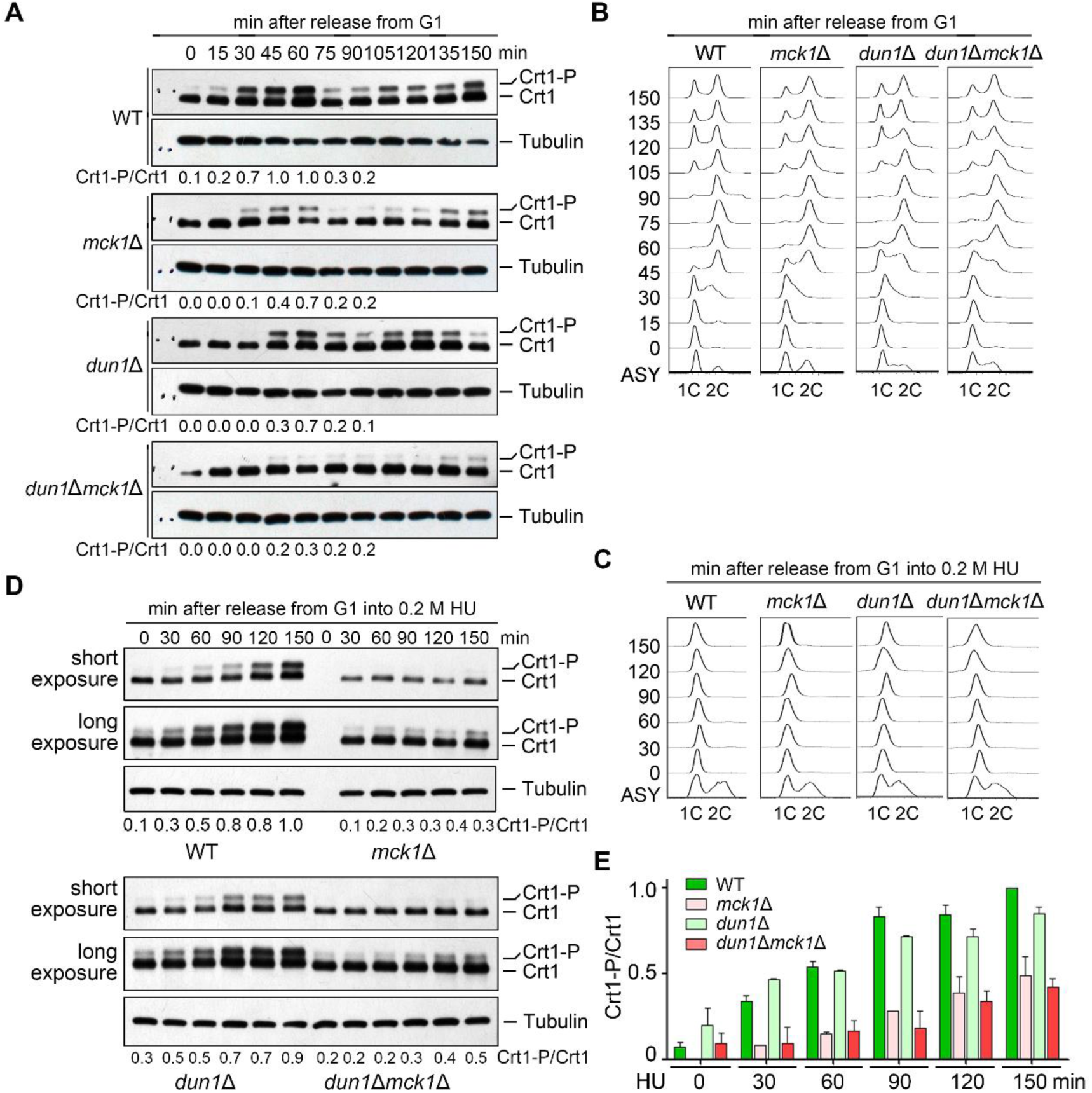
Mck1 is responsible for both cell-cycle-dependent and HU-induced Crt1 phosphorylation. (A) A representative cell cycle profile of the samples used for time course analysis of Crt1 phosphorylation in (B). (B) Phosphorylation of Crt1 in normal S phase. Cells were synchronized in G_1_ by α-factor before release into fresh media for the indicated times. The cell cycle progression was analyzed by FACS. Lysates were prepared from WT and *mck1*Δ cells. The protein levels of Crt1-13MYC were detected by immunoblots using an anti-MYC antibody as described as in Fig. 2D. The Crt1-P/Crt1 ratio was indicated below each lane for the first cell cycle. (C) A representative cell cycle profile of the indicated strains in the presence of 200 mM HU. (D) HU-induced Crt1 phosphorylation. Cells were synchronized in G_1_ by α-factor before release into fresh media containing 200 mM HU for the indicated times. The cell cycle progression was analyzed by FACS. Lysates were prepared from WT and *mck1*Δ cells. The protein levels of Crt1-13MYC were detected by immunoblots using an anti-MYC antibody as described as in Fig. 2D. See Fig. S3C for the results of biological repeats. (E) Quantitation of the relative levels of Crt1 phosphorylation from biological repeats.

In the presence of 0.2 M HU, the cell cycle progression of all alleles was almost completely halted within 150 min (Fig. 3C). This indicates an intact S phase arrest function of the replication checkpoint in these mutants, consistent with a role of Mck1 and Dun1 downstream of Rad53. Importantly, Crt1 phosphorylation occurred more slowly with a significant lower level in *mck1*Δ than in *dun1*Δ and WT (Fig. 3D, 3E and S3C). These data suggest a critical role of Mck1 in Crt1 phosphorylation in both normal and perturbed conditions.

### Phosphorylation-mimetic mutations of *CRT1* compensates the checkpoint defect of *mck1*Δ

To address the physiological significance of Mck1-mediated Crt1 phosphorylation, we reasoned that Mck1 may target Crt1 to antagonize its repressor function.

We first tested whether Crt1 phosphorylation can suppress the ultra-sensitivity of *dun1*Δ*mck1*Δ to HU as *crt1*Δ. Crt1 comprises nine putative Mck1 recognition motifs (S1-S9, Fig. 4A), (S/T)-x-x-x-(pS/T)*, where * stands for the priming phosphorylated residue and x for any amino acid [42]. We mutated these serine or threonine residues to aspartic acids to mimic the phosphorylation state. To examine the suppression effect of *crt1* mutations, the plasmids expressing various *crt1* alleles were transformed into the *dun1*Δ*mck1*Δ*crt1*Δ triple mutant. Consistently, a plasmid expressing WT *CRT1* prominently sensitized *dun1*Δ*mck1*Δ*crt1*Δ to 50 mM HU (Fig. 4B, compare lines 9 and 10). Through a series of different combinations, we found that phospho-mimetics of many sites (e.g., S222/T226 and S295/S299) are capable to suppress the HU sensitivity of the triple mutant to various extents (Fig. 4B, lines 3 and 13). These results suggest that these putative Mck1 sites play partially redundant roles in the S-phase checkpoint. Among them, *crt1-S5D (S295DS299D)* rescued the growth of the *dun1*Δ*mck1*Δ*crt1*Δ (Fig. 4B, compare line 13 to 9) triple and *mck1*Δ*crt1*Δ double mutants (Fig. S4) to an extent comparable to the empty vector, indicating that we have isolated a complete loss-of-function phospho-mimetic mutant of the Crt1 repressor. These results imply that Mck1 kinase abrogates the repressor function of Crt1 through phosphorylation (predominantly at the Mck1 kinase consensus sites S295/S299).

**Figure 4.**
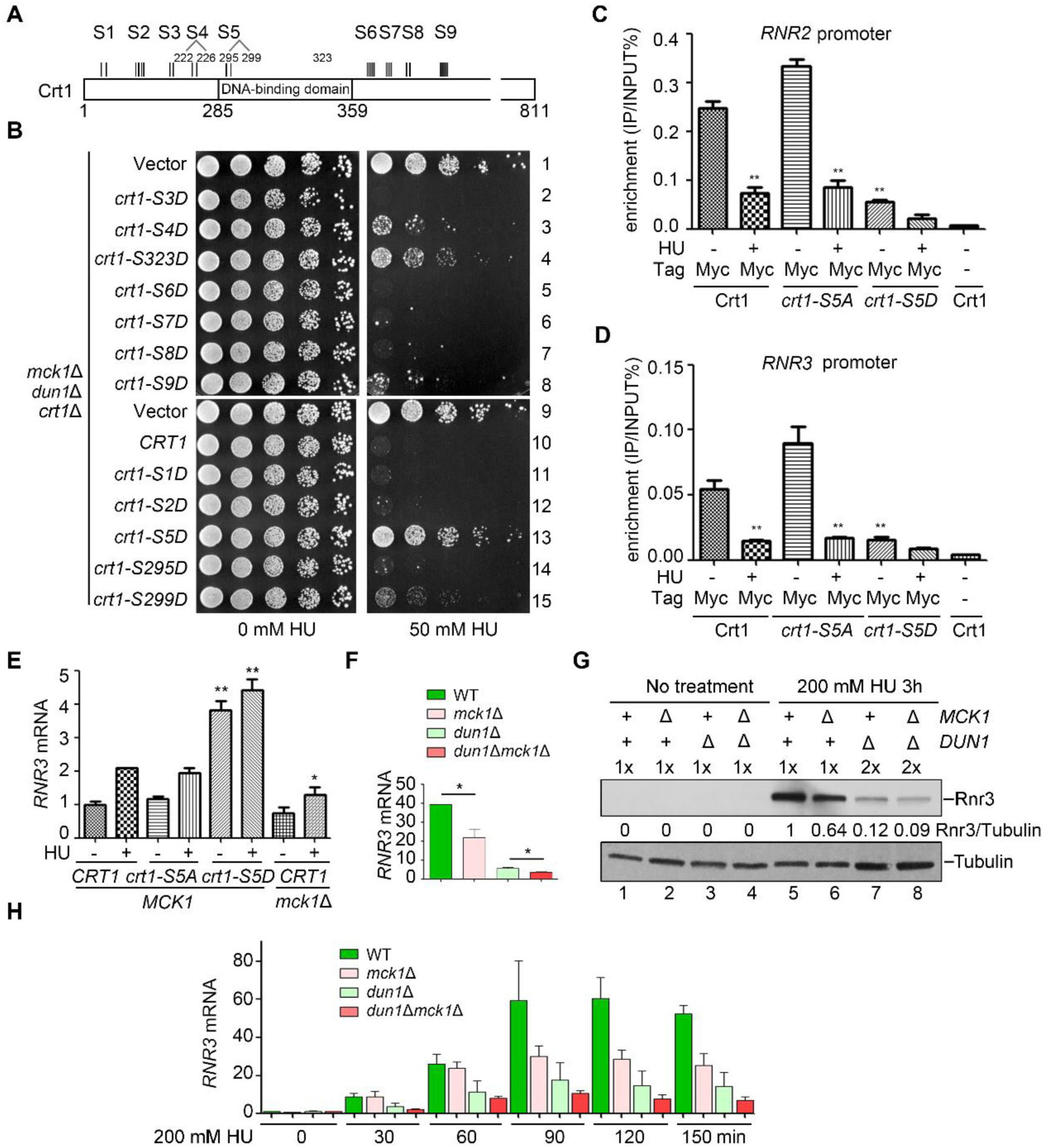
Crt1 phosphorylation affects its binding to the promoters of *RNR* genes and thereby reduces *RNR* induction. (A) A schematic diagram of the consensus Mck1 kinase recognition motifs in Crt1. Crt1 bears eight clusters of (S/T)-X-X-X-(pS/T)*, where * stands for the priming phosphorylated residue and X for any amino acid. S1 (S58, S62); S2 (S167, S171, S173, S174); S3 (T197, T199); S4 (S222, T226); S5 (S295, S299); S6 (S388, S389, S391, S393, S394); S7 (S412, S414, T416, S418); S8 (T488, S492); S9 (S556, S558, S560, S562). (B) Phospho-mimic mutations of several putative Mck1 sites in Crt1 rescue the HU sensitivity of *mck1*Δ*dun1*Δ to a similar extent as *crt1*Δ. The *dun1*Δ*mck1*Δ*crt1*Δ triple mutant was transformed with pRS313 vector, WT *CRT1* or the indicated mutants. Five-fold dilution of the cells were grown at 30°C for 48 h. (C) ChIP-qPCR assays of Crt1 localization at the *RNR2* promoter in various alleles in the presence or absence of 200 mM HU. (D)ChIP-qPCR assays of Crt1 localization at the *RNR3* promoter in various alleles in the presence or absence of 200 mM HU. (E, F) Measurement of the *RNR3* mRNA levels in various mutants by qPCR. Cells were grown in rich media with or without 200 mM HU for 3h. The total mRNA was prepared as Methods and Materials. The relative levels of *RNR3* to *ACT1* mRNAs were determined by qPCR analyses. The value of untreated WT was set to 1.0. Error bars represent standard deviations from at least three biological repeats. *P*-value <0.01 and 0.05 are donated as “**” and “*”, respectively. (G) Western blotting of the Rnr3 protein levels in various mutants with or without 200 mM HU treatment for 3 h. Mck1 is necessary for efficient Rnr3 induction. Cells were grown and treated as above. Rnr3-13MYC and Tubulin were detected by immunoblots. (H) Time course measurement of the *RNR3* mRNA levels by RT-PCR after HU treatment for the indicated time. Three biologically repeated experiments were basically performed as described in (E).

### Mck1-dependent phosphorylation of Crt1 abrogates its promoter binding

Interestingly, the dominant Mck1 sites S295 and S299 are located within the DNA binding domain of Crt1 (Fig. 4A), raising the possibility that phosphorylation of these sites may regulate its DNA binding capability. Therefore, we next examined the binding of Crt1 on the *RNR* promoters through chromatin immunoprecipitation (ChIP). Crt1 was significantly enriched at the promoter regions of both *RNR2* (Fig. 4C) and *RNR3* (Fig. 4D), which was dramatically reduced after 0.2 M HU treatment. These results indicate that Crt1 dissociates from the promoters of *RNR2* and *RNR3* in response to HU. Nevertheless, the phospho-mimetic mutant proteins (*crt1-S5D*) retained only approximately 20% enrichment of that of WT even in the absence of HU. Notably, Crt1-S5A still maintained the response to HU, indicating that Crt1 bears other Mck1 sites (e.g., S222 and S226) and/or phosphorylation sites of kinases other than Mck1 (e.g., Dun1 and Rad53). Taken together, these data suggest that Mck1-mediated Crt1 phosphorylation compromises the DNA binding activity and thereby regulates the repressor function of Crt1.

To directly test this, we next checked the expression of the Crt1-controlled genes. In *crt1-S5D*, *RNR3* was constitutively expressed in significantly higher level than in WT and *crt1-S5A* even in the absence of HU (Fig. 4E). This result is consistent with its suppression effects on *mck1*Δ shown in Fig. 4B and S4, indicating that phosphomimetic mutation of these Mck1 sites is sufficient to abrogate the repression by Crt1. Similarly, the induction of *RNR3* upon HU treatment was significantly compromised in *mck1*Δ though to a lesser extent than in *dun1*Δ at both transcriptional (Fig. 4E and 4F) and translational levels (Fig. 4G), which is congruent with their relative HU sensitivity (Fig. 2D). To further address the contribution of Mck1 and Dun1 in *RNR3* induction, we performed time course analysis of *RNR3* transcription. In the absence of HU, *RNR3* was barely expressed in G_1_-arrested cells (Fig. 4H). After release into 200 mM HU for 30 min, the *RNR3* transcripts were gradually elevated, which was prominently impaired in *dun1*Δ. Intriguingly, there was a stark rise in the *RNR3* mRNA levels around 90 min in WT cells, which was significantly compromised when *MCK1* was deleted. However, unlike *dun1*Δ, *mck1*Δ did not show an apparent effect during the initial induction stage (0-60 min), indicating that Mck1 likely functions kinetically later than Dun1 or through an indirect effect in response to HU (Fig. 4H). This is also in good agreement with the observations that *mck1*Δ does not display sick growth in the presence of the moderate concentrations of HU (Fig. 1 and 2). Taken together, these results argue for a critical role of Mck1 in antagonizing the Crt1 repression of *RNR* genes. These data also implicate that Dun1 acts as a primary kinase in initiating *RNR3* induction, while Mck1 might be required for the additional augment when more *RNR* expression is needed (e.g., severe and/or persistent stress).

### Mck1 also regulates the Rnr2 inhibitor Hug1

As shown in Fig. 2D, *CRT1* deletion only shows a partial suppression of the *mck1*Δ phenotype. This raises a possibility that there are additional Mck1 targets besides Crt1. To test this, we carefully compared the suppression effects of all known negative regulators of RNR.

Consistent with the results shown in Fig. 2A, if we removed only one of the RNR-hijacking proteins including Sml1, Hug1, Dif1 and Wtm1, there was no detectable effects in both *mck1*Δ (Fig. 5A and S5A) and *mck1*Δ*dun1*Δ mutants (Fig. S5B). The possible reasons are: 1) Mck1 does not mainly contribute to regulating the RNR protein localization and/or nuclear-cytoplasmic trafficking; 2) the effects of Mck1 in RNR post-translational regulation may be masked by its dominant effects on the *RNR* expression level.

**Figure 5.**
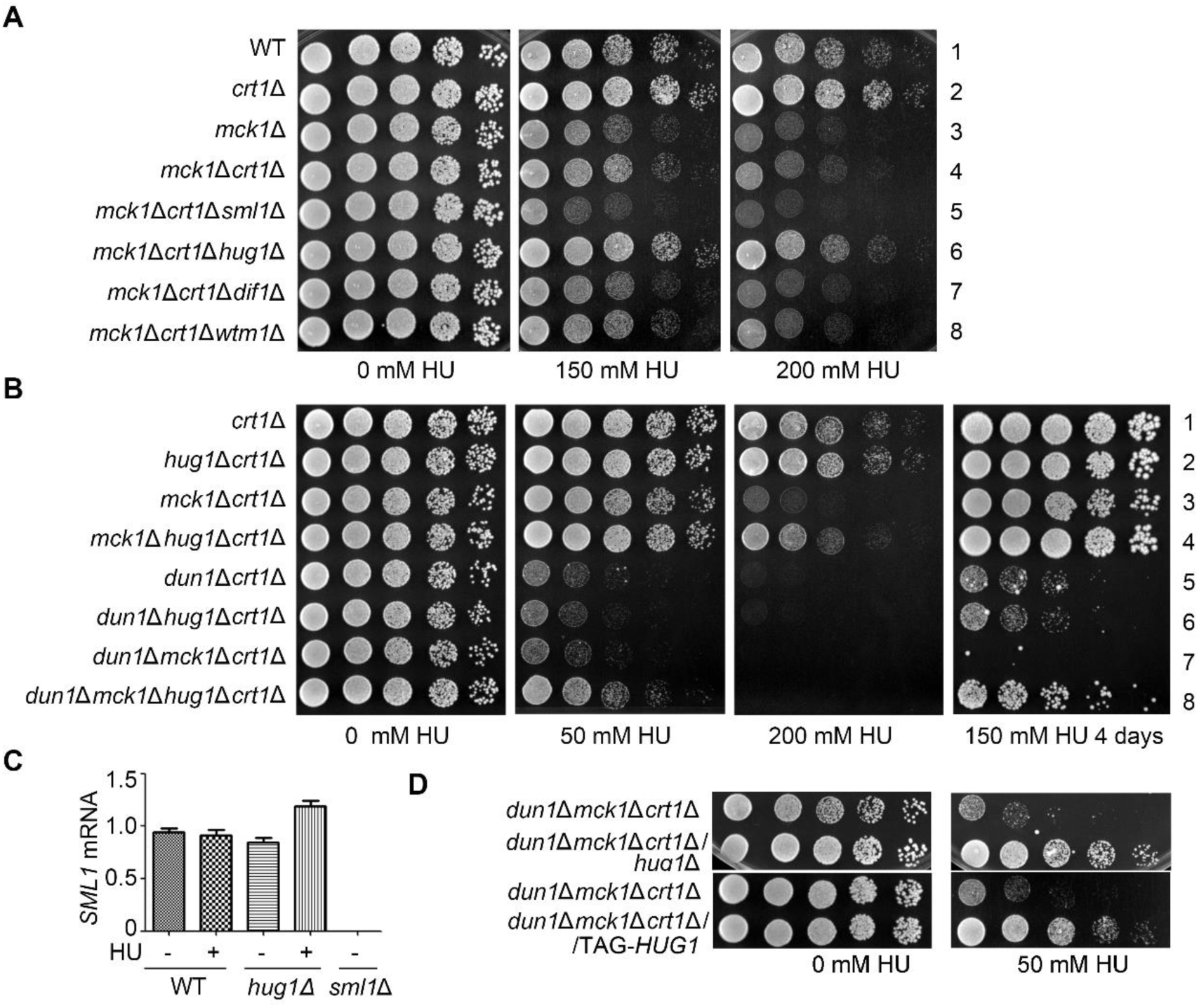
Hug1 suppresses the HU sensitivity of *mck1*Δ as a protein. (A, B) *HUG1* deletion, in combination with *CRT1* deletion, shows additive suppression effects on *mck1*Δ (A) and *mck1*Δ*dun1*Δ (B), but not on *dun1*Δ (B). Yeast strains with the indicated genotype (Table S1) were tested for the HU sensitivity by serial dilution analysis as described in Fig.1A. (C) *HUG1* deletion does not reduce *SML1* transcription. RT-PCR analysis of *SML1* mRNA in *hug1*Δ with or without HU treatment for 3 h. (D) Hug1 acts as a protein. A strain carrying TAG-*HUG1*, with its start codon (ATG) replaced by a stop codon (TAG) was tested in parallel with *hug1*Δ for the HU sensitivity in the indicated genotypes as described in (A).

To test these possibilities, we next eliminated each RNR-binding protein together with the major suppressor Crt1. Consistently, *CRT1* deletion showed suppression in either *mck1*Δ (Fig. 5A, line 4), *dun1*Δ (Fig. S5C, line 3) or *mck1*Δ*dun1*Δ (Fig. S5B, line 8). Further removal of Sml1, Dif1 or Wtm1 had no additive effects with *crt1*Δ on either *mck1*Δ (Fig. 5A, lines 5, 7 and 8; Fig. S5D) or *dun1*Δ (Fig. S5C, lines 4-7). When we deleted *CRT1* and *HUG1* in combination, we found a synergistic rescue on *mck1*Δ (Fig. 5A, compare line 6 to 4) and *mck1*Δ*dun1*Δ (Fig. 5B, compare line 8 to 7). On the contrary, *hug1*Δ*crt1*Δ exhibited no more suppression for *dun1*Δ than *crt1*Δ alone (Fig. 5B, compare line 6 to 5). Although *HUG1* locates adjacent to *SML1* on the genome, *HUG1* deletion did not reduce *SML1* transcription (Fig. 5C). Moreover, despite the short length of *HUG1* gene (207 bp), it indeed acted as a protein because a nonsense mutation of the sole start codon (ATG replaced by TAG) led to the same suppression as *hug1*Δ (Fig. 5D). These results suggest that besides Crt1, Hug1 protein is an additional key downstream effector of Mck1.

To confirm this, we then examined the Hug1 protein levels in these mutants. Under normal condition, Hug1 protein was barely detectable in WT (Fig. 6A, lane 1). Knockout of *MCK1*, but not *DUN1*, elevated the Hug1 levels, though to a lesser extent than *crt1*Δ (compare lanes 2, 3 and 9). Apart from the previously reported repression by Crt1 at the transcriptional level [18], these results indicate that *HUG1* is also repressed by Mck1 kinase in normal condition. Under 200 mM HU treatment, *mck1*Δ caused a prominent increase comparable to *crt1*Δ, whereas the *mck1*Δ*crt1*Δ double mutant resulted in a synergistic augment of the Hug1 level (Fig. 6A, compare lanes 6, 13 and 14). On the contrary, deletion *DUN1* antagonized the *HUG1* induction caused by *crt1*Δ (compare lanes 13 and 15) and *mck1*Δ (compare lanes 6, 8, 14 and 16). These data suggest that Mck1 regulates *HUG1* expression in a Crt1-independent manner.

**Figure 6.**
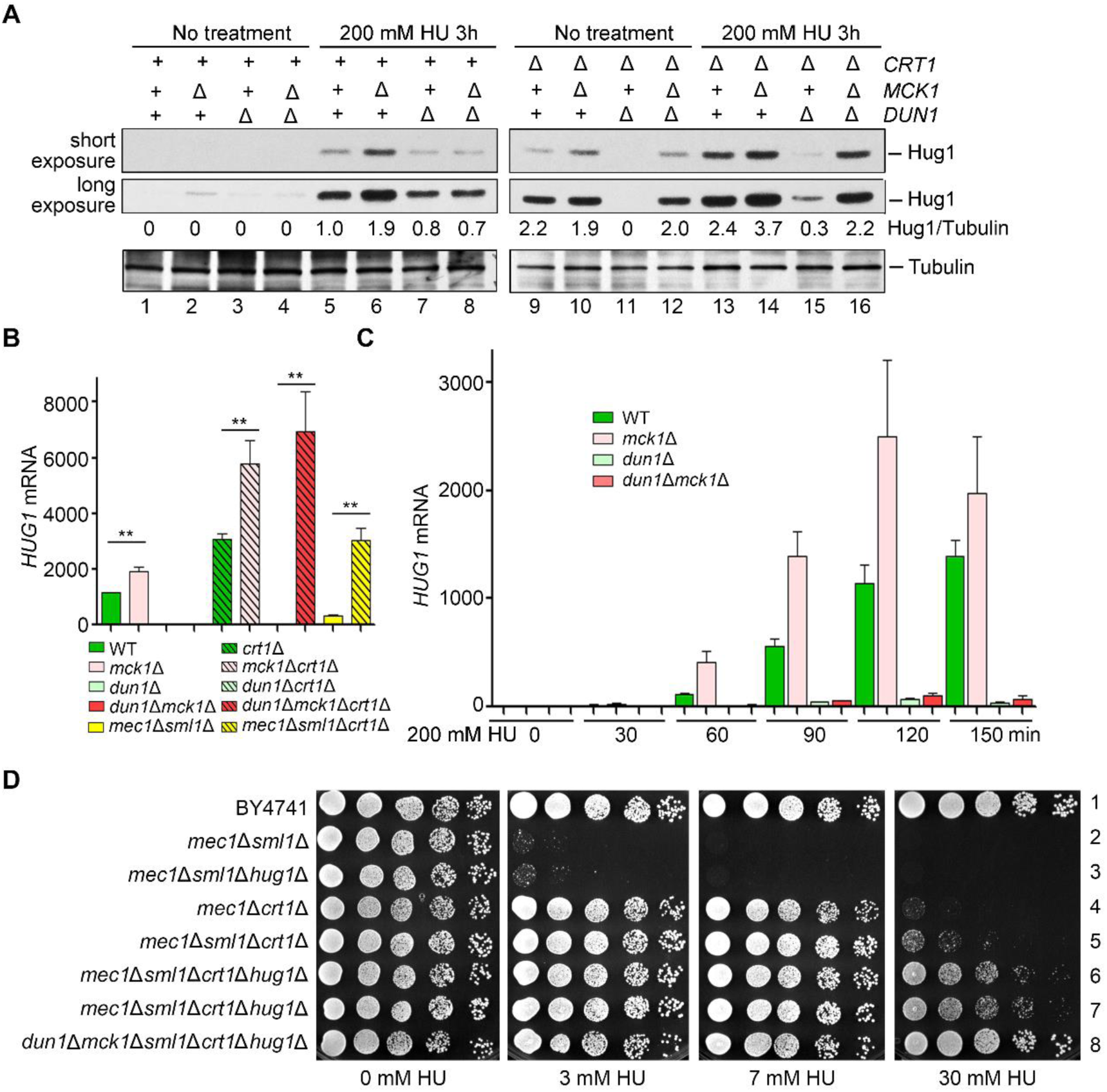
Mck1 regulates the transcription of *HUG1* in a Crt1-independent way. (A) Mck1 and Crt1 co-regulate the Hug1 protein level. Cells were grown and treated as Fig. 4C. Hug1-13MYC and Tubulin were immunoblotted with anti-MYC and anti-tubulin antibodies, respectively. (B, C) RT-PCR analysis of the *HUG1* mRNA levels relative to *ACT* in various alleles after HU treatment for 3 h (D) or the indicated time points (E). The value of untreated WT was set to 1.0. (D) Combinational deletion of *SML1*, *CRT1* and *HUG1* displays additive suppression effects on *mec1*Δ and *mck1*Δ*dun1*Δ. Yeast strains with the indicated genotype (Table S1) were tested for the HU sensitivity by serial dilution analysis as described in (A).

### Mck1 negatively controls Hug1 at the transcriptional level

Next, we asked how Mck1 regulates the cellular levels of Hug1. Hug1 is unlikely a substrate of Mck1 kinase since there are no Mck1 consensus recognition sites. Thus, we examined the *HUG1* mRNA levels after 200 mM HU treatment for 3 h. Consistently, deletion of *MCK1* and *CRT1* individually led to an increase of nearly 100% and 150% in the *HUG1* mRNA levels compared to WT, respectively, whereas deletion of *MCK1* and *CRT1* in combination resulted in an approximately 400% increase (Fig. 6B). On the other hand, *dun1*Δ eliminated induced *HUG1* transcription in *crt1*Δ and *mck1*Δ, but not in *crt1*Δ*mck1*Δ double mutant. Further removal of *CRT1* led to a maximum *HUG1* induction, confirming that *HUG1* is repressed by Crt1 which is relieved by Dun1. In good agreement with the protein levels of Hug1 mentioned above, these data allow us to conclude that Mck1 inhibits the *HUG1* induction at the transcriptional level.

Next, we quantitated the *HUG1* mRNA levels at 30-min intervals after 0.2 M HU treatment. Transcription of *HUG1* was elevated more than 1000-fold within 2 h after HU treatment (Fig. 6C). In the absence of *MCK1*, the induction of *HUG1* nearly doubled than WT at each time point. Putting together, these data suggest that *HUG1* transcription is exquisitely controlled by a pair of antagonistic mechanisms of the S-phase checkpoint, induction by elimination of the Crt1 repressor function (mainly through Dun1 kinase) and direct inhibition by Mck1 kinase.

Since *mck1*Δ*dun1*Δ was identified to mimic *mec1*Δ or *rad53*Δ in response to HU (Fig. 1B), we then tested whether deletion of the main targets of Mck1 and Dun1 is able to suppress the HU sensitivity of their upstream kinase mutants as well. Deletion of *CRT1* alone was sufficient to afford *mec1*Δ to resist 7 mM HU, whereas *SML1* deletion could not (Fig. 6D, compare lines 2, 3 and 4). Combined deletion of *CRT1* and *SML1* slightly facilitated HU resistance of *mec1*Δ (line 5). We further deleted *HUG1*, and found significant enhanced HU resistance in *mec1*Δ (Fig. 6D, lines 6 and7) as well as *mck1*Δ*dun1*Δ double mutant (line 8). These data suggest that Crt1, Hug1 and Sml1 represent the major effectors of the Mec1-Rad53-Dun1/Mck1 kinase cascade in RNR regulation.

### Mck1 directly participates in the stress-induced dNTP regulation

We next directly compared intracellular dNTP pools in WT or *mck1*Δ-related mutants. Because the dNTP levels are cell-cycle controlled, cells were first arrested in G_1_ before release into S phase in the presence of 200 mM HU. Considering that Mck1 may function kinetically later as shown in Fig. 4H, we collected the cells after HU treatment for 0, 3 and 6 h for dNTP measurement. In the G_1_ cells before HU assault (0 h), all strains carrying *mck1*Δ had a moderately increase in dNTP pools compared with WT (Fig. 7A), suggesting a role of Mck1 in regulating the dNTPs levels and balance in normal condition. Chronic HU treatment elicited a dramatic decrease of dNTP pools, with the highest decrease in dATP, which thus became the most limiting of the four dNTPs instead of dGTP in the normal condition (Fig. 7B and 7C) (Kumar et al, 2010). These indicate that HU causes dNTP imbalance as well as depletion. In WT and *dun1*Δ cells, dNTP levels were partially restored in 6 h. However, dNTP restoration was abolished in *mck1*Δ and *mck1*Δ*dun1*Δ. Strikingly, dATP remained extremely low in both mutants. These results indicate that the recovery of dNTP homeostasis (including both levels and balance) is dependent on Mck1 in the presence of high concentration HU assault.

**Figure 7.**
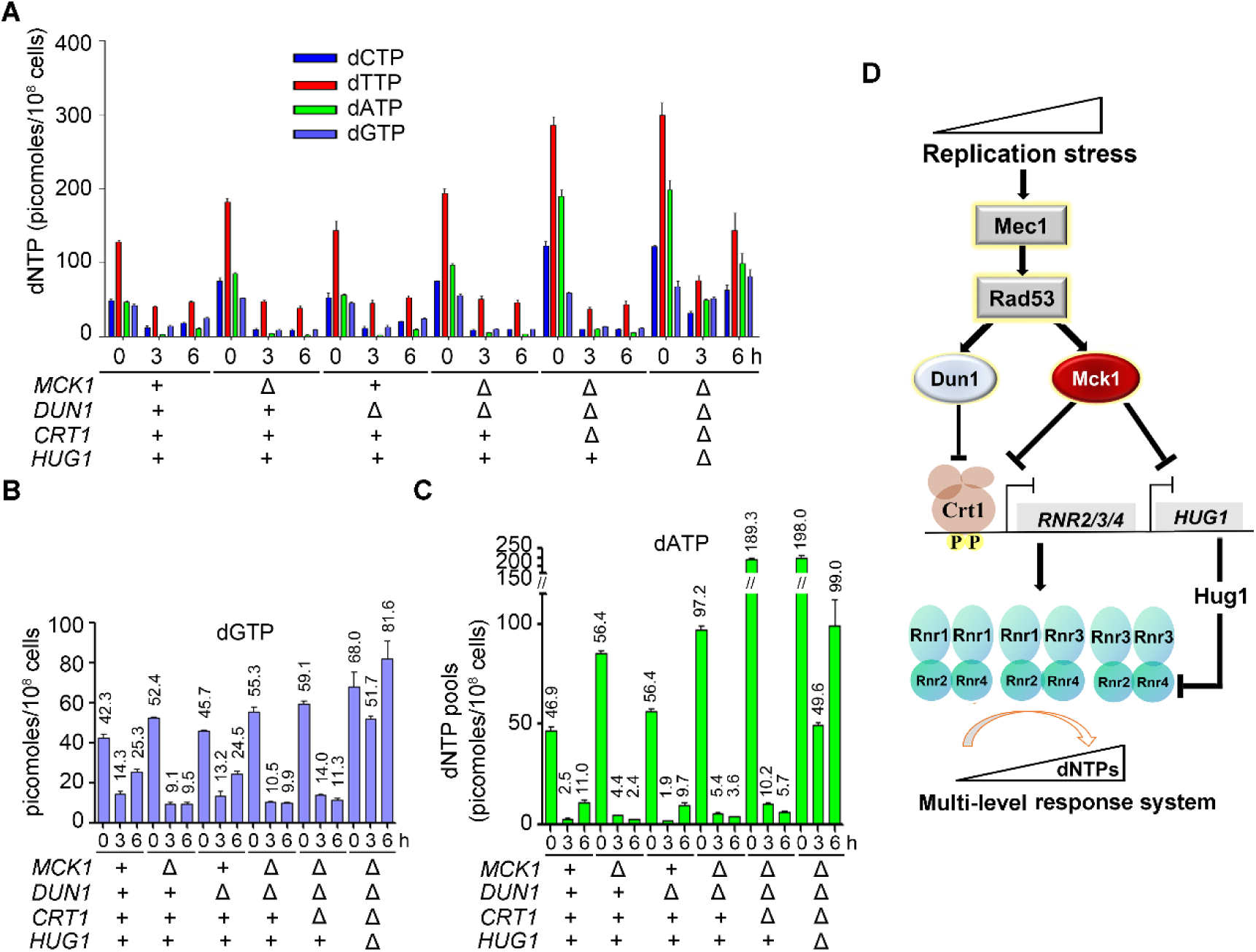
Mck1 regulates cellular dNTP pools in response to a lethal dose of HU. (A, B, C) dNTP (A), dGTP (B) or dATP (C) levels correlate with HU-resistant phenotypes of the mutants. dNTPs were measured for cell cultures of the indicated genotypes. Cells were arrested in G_1_ and then released into 200 mM HU for 0, 3 and 6 h. dNTP levels were normalized to NTP levels in each sample, and then divided by total number of cells used for the preparation. Error bars reflect standard deviation (SD) of the three biological repeats. (D) A proposed model for the roles of Mck1 as the downstream kinase of Mec1-Rad53 in response to replication stress. Under perturbed condition, Mck1, together with Dun1, antagonizes the repressor function of Crt1 via phosphorylation, which in turn allows the derepression of *RNR2/3/4* transcription. Meanwhile, Mck1 inhibits the expression of Rnr2-hijacking protein Hug1, which is concomitantly induced due to the elimination of the Crt1 repressor. Therefore, Mck1 defines a Dun1-independent pathway in fine-tuning RNR activity to maintain appropriate dNTP pools according to the degrees of replication threats. For simplicity, the effectors known to be targeted mainly by Dun1 (i.e., Sml1, Dif1, Wtm1) are not shown here.

*CRT1* deletion alone led to only a mild elevation of the dNTP levels in the presence of 200 mM HU (Fig. 7B). Strikingly, when *HUG1* was further deleted, we observed a dramatic expansion of dNTP pools, congruent with the percentages of *HUG1* induction in the *mck1*Δ*dun1*Δ*crt1*Δ triple mutant shown in Fig. 6. Among them, dATP was augmented most significantly, suggesting that Hug1 is also able to restore the balance of four dNTPs impaired by HU. These results indicate that Hug1 is a potent suppressor of the RNR activity, particularly when Crt1 repressor function is abrogated. Importantly, the recovery of dNTP levels correlated with the growth of these mutants in the presence of HU (Fig. 5A, 5B and 6D). Putting together, we propose that Mck1 defines a secondary effector branch of the Mec1-Rad53 cascade and plays a crucial role for cells coping with a more severe and/or long-lasting replication insult (Fig. 7D).

## Discussion

Dun1-independent *RNR* induction in response to either exogenous or endogenous replication stress have been observed by different groups [18, 43]. Nearly two decades after the initial report, here we have identified that Mck1 is a new downstream kinase of Rad53 and functions in the Dun1-independent pathway in dNTP regulation.

Cells need to maintain the appropriate amount and balance of all four dNTPs, an even more challenging task when they suffer exogenous or endogenous replication stress. Moreover, cells should have multi-layer response systems to deal with various degrees of stress. Here we prove that Mck1 and Dun1 kinases cooperate with each other to achieve this. Under unperturbed condition, Crt1 represses the expression of *RNR* genes to avoid overproducing dNTPs. Under moderate perturbed condition, the Mec1-Rad53 cascade activates Mck1 and Dun1. At the post-translational level, Dun1 is responsible for releasing the caged Rnr1 (by Sml1) and Rnr2/4 (by Dif1 and Wtm1), which allows more RNR holo-enzyme formation. At the transcriptional level, Dun1 and Mck1 alleviate the repressor function of Crt1 through phosphorylation at different sites with different kinetics, allowing a gradual de-repression of *RNR2/3/4*. Meanwhile, Crt1-controled *HUG1* is also induced, which very likely prevents overproducing dNTPs under this condition. Excessive dNTPs have been demonstrated to have an effect on increasing mutation risk and thus impair cell growth [43, 44]. However, the higher levels of RNR activity may be required to produce enough dNTPs if cells suffer a more severe and/or persistent assault (i.e. more than 150 mM HU). Mck1 operates under this circumstance through inhibiting the induction of *HUG1* in a Crt1-independent manner. Apart from the dNTP levels, the Mck1-Hug1 pathway also regulates the dNTP balance under replication stress induced by HU. Although molecular details regarding how Mck1 and Hug1 achieve these need further investigation, our findings reveal a multi-level response system to a wide range of replication threats.

It is also noteworthy to point out that the low dNTP levels are unlikely the sole reason underlying the high HU sensitivity of *mck1*Δ*dun1*Δ. Therefore, it will be interesting to search for additional roles of Mck1 in maintaining genome stability other than the mechanism reported here. Intriguingly, apart from the RNR regulation function reported here, Mck1/GSK-3 has been well-established as phosphodegrons of an array of vast substrates including cell cycle proteins like Cdc6 [45], Sld2 [46], Hst3 [47], Eco1 [48] in yeasts, and Bcl3, c-JUN, Mdm2, c-Myc, Rb and PTEN in mammals [49], which are all important for cell growth and proliferation.

Although there are no apparent orthologs of Hug1 in higher eukaryotes, several other studies have provided some hints that the role of Mck1/GSK-3 in the S phase checkpoint might be conserved. An unusual feature of GSK-3 is that it is generally active under unperturbed condition and primarily regulated by inhibition in response to extracellular signals (e.g. growth factors, insulin) through signaling pathways like Akt and mTOR (Target Of Rapamycin)[50]. The TOR kinase, which belongs to the highly conserved family of phosphatidylinositol-3-kinase-related kinases (PIKKs) as Mec1^ATR^, has been shown to be required for DNA damage-induced expression of *RNR1* and *RNR3* in yeasts [51, 52]. In mammalian cells, the translation of large and small RNR subunits RRM1 and RRM2 is cap-dependent, which is regulated by phosphorylation of eukaryotic translation initiation factor 4E (EIF4E)-binding protein 1 (4E-BP1) by mTORC1 [53].

Thus, further investigation of the role of GSK-3 in the S-phase checkpoint and RNR regulation in vertebrates may help to establish crosstalk among glucose metabolism, DNA metabolism and cell proliferation. In consideration of the clinical usage of HU and pharmaceutical interest in the inhibitors of the cell cycle checkpoint proteins including Gsk-3 kinases for neoplastic and non-neoplastic disease treatments [7, 54–56], the studies based on our results reported here may have potential implications for drug design.

## Materials and Methods

### Yeast strains and plasmids

*S. cerevisiae* strains congenic with BY4741/4742 and plasmids constructed in this study are listed in Table S1 and Table S2, respectively.

### Synthetic genetic array (SGA)

The *dun1*Δ (MATα) single mutant was crossed with a non-essential deletion collection of cell cycle-related genes for synthetic genetic screens as previously described [57, 58]. The obtained double mutant colonies were then examined for their growth in the presence or absence of 15 mM HU.

### HU sensitivity assay (Spot assay)

Fivefold serial dilution of log phase growing cells (initial OD_600_ = 0.4) were spotted on YPD (yeast extract/peptone/dextrose) or synthetic media plates in the presence of indicated concentrations of HU. Plates were incubated at 30°C for 48 h before photography.

### *In Vitro* Kinase Assay

To expressed His6-Rad53 and His6-rad53-KD, the pET-15b-*RAD53* (a kind gift from Dr. John Diffley) and pET-15b-*rad53-KD* (K227A) plasmids were introduced in BL21-codon-plus(DE3)-RIL E. coli strain (Stratagene). Early log phase culture was treated with 0.2 mM IPTG to induce the protein expression. After 3 h of incubation at 25°C, cells were harvested. The proteins were purified using Ni^2+^-beads (GE Healthcare) and eluted by 250 mM imidazole. pRS-313-pADH1-*MCK1-5FLAG* plasmid was transformed into *mck1*Δ strain. Mck1-5FLAG was purified by 20 μl anti-FLAG M2 beads (Sigma) and eluted by 150 μl of 1 μg/μl FLAG peptide. In a typical kinase assay, 50 mM Tris, pH 7.5, 150 mM NaCl, 0.1% Tween-20, 10 mM MgCl_2_, 5 μCi of ɣ-^32^P-ATP were used. Each kinase reaction contained His6-Rad53 (0-10 μg) or His6-Rad53-KD (10 μg) and Mck1-5FLAG (10 μl) in 40 μl reaction volume and incubated for 30 min at 30°C. Kinase assay was stopped by heating at 100°C for 5 min in SDS sample buffer. Samples were then subject to SDS–PAGE. Phosphorylation was detected by ^32^P autoradiography. The amount of proteins loaded was detected by CBB staining.

### Protein detection

For immunoblot anaysis, 5 ml of culture was grown in YPD to an OD_600_ of 1 and harvested. Indicated culture was treated with 200 mM HU for 3h before being harvested. Yeast extracts were prepared using the trichloroacetic acid (TCA) precipitation for analysis in SDS-gels. The Crt1-13MYC, Rnr3-13MYC, Hug1-13MYC protein levels were detected with mouse anti-MYC antibody (1:1000, ORIGENE) and HRP-conjugated anti-mouse IgG as the secondary antibody (1:10000, Sigma). Tubulin as loading control was detected with anti-tubulin (1:10000, MBL) and HRP-conjugated anti-rabbit IgG as the secondary antibody (1:10000, Sigma). For detecting the heperphosphorylation of Crt1-13MYC, the special 30% Acrylamide Solution (acrylamide: N’N’-bis-methylene-acrylamide = 149:1) was used.

### Pulsed field gel electrophoresis (PFGE) and southern blot

Stationary phase cells (2.5 × 10^7^) were washed and re-suspended in 50 µl of Lyticase buffer (10 mM Phosphase buffer pH 7.0, 50 mM EDTA), and then solidified in blocks with 50 µl 1% low melting temperature agarose (Sigma). These were digested with 75 U/ml lyticase in Lyticase buffer for 24 h at 37°C, then with 2 mg/ml Proteinase K (Amresco) in 100 mM EDTA, 1% sodium lauryl sarcosine for 48 h at 42°C. After four washes with TE50 (10 mM Tris, pH7.0, 50 mM EDTA), plugs were run on 1% agarose gels on in 1× TBE at 3 V/cm, 300-900 s switch time, for 68 h. PFGE was carried out in a CHEF-MAPPER system (BioRad) for 68 h at 14°C. Chromosomes were visualized with ethidium bromide prior to treatment with 0.25 M HCl for 20 min, water for 5 min twice. DNA was transferred to HyBond N^+^ in transfer buffer (0.4 M NaOH, 1 M NaCl) and UV cross-linked before hybridization with a random primed probe (Takara) overnight at 42°C and washed twice for 20 min with 0.5× SSC 0.1% SDS at 65°C.

### Quantitative RT-PCR

Total RNAs extraction was performed using a commercial TRIzon Reagent (CoWin Biosciences) and the manufacturer’s instructions with slight modifications. After centrifugation, cells were added to 100 μl TRIzon Reagent together with 100 μl of sterile glass beads (0.5 mm in diameter). The cells were then disrupted by vertexing for 60 s followed by cooling on ice for 60 s. This step was repeated four times. The extraction was then continued according to the manufacturer’s instructions (CoWin Biosciences). For reverse transcription-PCR (RT-PCR) analysis, reverse transcription with Oligo (dT) Primer was performed with 2 μg of total RNA, 1 mM dNTPs, 1μl RT and 0.5 μl RNasin for 60 min at 42°C, which was followed by a 15 min heat inactivation at 95°C. For each gene, real-time quantitative PCR amplification (95°C for 10 min followed by 95°C for 15 s and 60°C for 1 min for 40 cycles) was performed using SYPR-Green on a QuantStudio 6 Flex system (Life).

### Chromatin Immunoprecipitation (ChIP)

Logarithmically growing cells were treated with formaldehyde prior to lysis. ChIP was carried out according to the methods used in previous studies with slight modifications. In brief, 100 ml stationary phase cells were treated with or without 0.2 M HU for 1 hr at 30°C. 1% formaldehyde was used for crosslinking for 20 min at room temperature. Cells were lysated and sonicated. Endogenous Crt1 proteins carrying a Myc13 tag were precipitated by an anti-Myc antibody (9E10) overnight at 4°C. The immune complexes were harvested by the addition of 50 µl of protein G dynabeads. Formaldehyde crosslinks were reversed by incubation at 65°C for 5 hr, followed by protease K treatment at 42°C for 2 hr. Then co-precipitated genomic DNA was purified using a phenol-chloroform extraction and subjected to quantitative real-time PCR SYPR-Green on a QuantStudio 6 Flex system (Life).

### dNTP Measurement

dNTP extraction and quantification were carried out as described [59].

## Supporting information

Supporting information (Tables S1-S2, Figures S1-S5) are available online.

## Author contributions

X.Li. performed most of the experiments; X.J and S.S. measured dNTP pools; X.Liu, J.Z. and Q.C. helped in strain/plasmid constructions, phenotyping and phosphorylation assays; Y.N. and J.L. initiated this study and identified the synthetic HU sensitivity between *mck1*Δ and *dun1*Δ; Z.L. carried out the mass spectrometry analysis. H.L., X.Li. and B.L designed the experiments and wrote the paper.

## Acknowledgments

We thank Dr. Andrei Chabes for helping with dNTP pool analysis and critical comments, Dr. John Diffley for sharing plasmids, Drs. Li-Lin Du, Judith L. Campbell, Cong Liu and members of the Lou lab for helpful discussion.

## Supporting information captions

**S1 Fig (Related to Fig 1). HU sensitivity analysis of Gsk-3 homologs and their double mutants**

Yeast strains with the indicated genotype (Table S1) were tested for the HU sensitivity by serial dilution analysis as described in Fig.1A.

**S2 Fig (Related to Fig 1). Mck1 shows synthetic interactions with checkpoint factors in response to HU.**

(A, B) Yeast strains with the indicated genotype (Table S1) were tested for the HU sensitivity by serial dilution analysis as described in Fig.1A.

(C) Representative tetrad dissection analyzed using the diploid cells with the indicated genotype.

**S3 Fig (Related to Fig 3). Mck1 acts as a kinase in response to HU.**

(A) Mck1 acts as a kinase in response to HU. The *dun1*Δ*mck1*Δ strain was transformed with pRS316 empty vector, WT *MCK1* or *mck1* alleles (the catalytic mutant allele, D164A; the activation-loop mutant allele, Y199F). Strains were spotted onto SC-Ura media with or without 50 mM HU and grown at 30°C for 48 h.

(B) Mck1 affects Crt1 phosphorylation. Cells were grown to stationary phase. Lysates were prepared and resolved by a 7% polyacrylamide (acrylamide: N’N’-bis-methylene-acrylamide = 149:1) gel containing SDS. The phosphorylation of Crt1-13MYC were detected by immunoblots using an anti-MYC antibody. Tubulin was applied as a loading control.

(C) Biological repeats of Figure 3D.

**S4 Fig (Related to Fig 4). HU sensitivity analysis of the suppression effects of *crt1-5D* mutant.**

Yeast strains with the indicated genotype (Table S1) were tested for the HU sensitivity by two-fold serial dilution analysis.

**S5 Fig (Related to Fig 5). HU sensitivity analysis of the suppression effects on various mutants.**

Yeast strains with the indicated genotype (Table S1) were tested for the HU sensitivity by serial dilution analysis as described in Fig.1A.

